# Proteomic Analysis Defines Kinase Taxonomies Specific for Subtypes of Breast Cancer

**DOI:** 10.1101/122739

**Authors:** Kyla. A. L. Collins, Timothy. J. Stuhlmiller, Jon. S. Zawistowski, Michael. P. East, Trang T. Pham, Claire R. Hall, Daniel R. Goulet, Samantha M. Bevill, Steven P. Angus, Sara H. Velarde, Noah Sciaky, Tudor I. Oprea, Lee M. Graves, Gary L. Johnson, Shawn M. Gomez

**Affiliations:** Curriculum in Bioinformatics and Computational Biology, University of North Carolina at Chapel Hill, Chapel Hill, NC 27514.; Department of Pharmacology, University of North Carolina at Chapel Hill, Chapel Hill, NC 27514.; Lineberger Comprehensive Cancer Center, University of North Carolina at Chapel Hill, Chapel Hill, NC 27514.; Joint Department of Biomedical Engineering, University of North Carolina at Chapel Hill and North Carolina State University, Chapel Hill, NC 27514.; Translational Informatics Division, School of Medicine, University of New Mexico, Albuquerque, NM 87131.

**Author notes:** These authors contributed equally to this work (Kyla. A. L. Collins, Timothy. J. Stuhlmiller).

## Abstract

Multiplexed small molecule inhibitors covalently bound to Sepharose beads (MIBs) were used to capture functional kinases in luminal, HER2-enriched and triple negative, basal-like and claudin-low breast cancer cell lines and tumors. Kinase MIB-binding profiles at baseline without perturbation proteomically distinguished the four breast cancer subtypes. Kinases lacking defined functions in breast cancer were highly represented in the MIB-binding taxonomies. We show that these understudied kinases, whose disease associations and pharmacology are generally unexplored, are integrated in kinase signaling subnetworks with kinases that have been previously well characterized in breast cancer. Computationally it was possible to define subtypes using profiles of less than 50 of the more than 300 kinases bound to MIBs that included understudied as well as metabolic and lipid kinases. Furthermore, analysis of MIB-binding profiles established potential functional annotations for these understudied kinases. Thus, comprehensive MIBs-based capture of kinases provides a unique proteomics-based method for integration of poorly characterized kinases of the understudied kinome into functional subnetworks in breast cancer cells and tumors that is not possible using genomic strategies. The MIB-binding profiles readily defined subtype-selective differential adaptive kinome reprogramming in response to targeted kinase inhibition, demonstrating how MIB profiles can be used in determining dynamic kinome changes that result in subtype selective phenotypic state changes.

## Introduction

In 2014 the NIH established an initiative entitled *Illuminating the Druggable Genome (IDG)* to determine the function of understudied proteins including kinases encoded in the human genome (https://commonfund.nih.gov/idg/index). The human kinome is comprised of ~520 protein kinases that are highly druggable using both competitive small molecule and allosteric inhibitors. Including both lipid and metabolic kinases enlarges this family to ~634 druggable kinases. Of the protein kinases, the function of about one-third are poorly defined with the function and regulation of 50-100 kinases remaining largely unknown. To categorize our understanding of proteins in the human genome the *IDG Knowledge Management Center (KMC)* (http://targetcentral.ws/index) has developed a set of criteria for target development level (TDL) for druggable proteins such as kinases, G protein-coupled receptors and ion channels [1]. This knowledge base was used to categorize the 634 kinases in the human genome as Tclin (50 kinases), Tchem (390 kinases), Tbio (163 kinases) and Tdark (31 kinases). Using TDL criteria, the KMC defines Tclin as *bona fide* disease-involved kinases that are drug targets for at least one FDA approved pharmaceutical entity. Tchem includes target kinases having characterized small molecules that bind with high potency (K_d_ ≤ 30nM), have active pharmacologic studies in relation to a disease, and there are often medicinal chemistry efforts to identify highly selective molecules to regulate function of the kinase. Tbio is a more difficult categorization of kinases and basically includes kinases that do not meet Tclin and Tchem standards. They may have an association with human disease and even small molecules that bind with high affinity. Finally, Tdark includes kinases having the least understanding of function and often there are few molecular tools available for their study. Tdark kinases are generally poorly characterized for their integration into kinase signaling networks, represent unknowns in disease associations, and are unexplored as selective drug targets alone or in combination with other drug targets.

Even with the growing databases of genomic information for different cancers, it is often still unclear how molecular taxonomies translate to phenotype. Additional methods characterizing proteomic taxonomies are needed to understand signaling networks, particularly of protein kinases due to their druggability. Important for this analysis of the cancer kinome is a characterization of understudied kinases, representing a third of the kinome and lacking essential functional characterization as well as molecular tools for their manipulation and study [2]. These understudied kinases need to be functionally integrated into kinase networks for a global understanding of kinome dynamics to be achieved both at baseline and in response to perturbation.

We focused on breast cancer as a model to address the integration of understudied kinases within kinase networks, which has three primary subtypes that include luminal (further sub-divided into luminal A and B subtypes) as well as the majority of HER2+ breast cancers along with triple negative breast cancer (TNBC), that can itself be broken into basal-like and claudin-low subtypes [3]. Interestingly, basal-like breast cancer using molecular taxonomies is as different from luminal and HER2+ breast cancers as lung cancer, leading to the proposal that basal-like breast cancers are in fact a unique disease [4,5]. Estrogen and progesterone receptor dependence and HER2 addiction define vulnerabilities in luminal/HER2+ breast cancers. However, in basal-like and claudin-low triple negative breast cancer, there are no oncogenic drivers that define a common vulnerability that can be therapeutically targeted.

In an attempt at having a more complete understanding of the integrated kinome in breast cancer, we have developed methods using multiplexed inhibitor beads (MIBs) coupled with mass spectrometry (MIB/MS) that have the ability to bind and identify a large percentage of kinases in the human kinome [6,7]. By RNA-seq, most cell lines express ~350 kinases and our MIB-binding profiling captures a significant percentage of the expressed kinome [8]. In the current study we have compiled the baseline kinase MIB-binding profile using MIB/MS for 15 cell lines across breast cancer subtypes in addition to patient tumors. It was possible to define kinase taxonomies for breast cancers using feature selection methodologies based on the MIB/MS profile of 50 kinases among the kinases captured by MIB/MS that includes understudied protein kinases, lipid and metabolic kinases. Using the baseline MIB-binding state in a machine-learning framework further allows accurate classification of breast cancer in both cell lines and primary tumors. Kinases identified within these distinguishing profiles are distributed throughout subfamilies of the kinome, representing multiple subnetworks with a significant representation of understudied kinases. In particular, we utilize a regression approach to integrate known interaction and phosphorylation data with MIB-binding behavior to establish functional subnetworks and associated annotation for 89 understudied kinases, including 22 kinases defined as Tbio or Tdark. These findings demonstrate that determining the functional kinome based on MIB-binding has prognostic value in defining the integration of signaling networks that is not currently possible using genomic strategies.

## Results

### Multiplexed Kinase Inhibitor Beads capture kinases from every subfamily and provide a means to assay understudied kinases

Multiplexed kinase inhibitor beads (MIBs) are a set of Sepharose beads each with a specific covalently-attached kinase inhibitor [6,9]. Coupling MIB gravity-flow affinity chromatography with mass spectrometry (MIB/MS) provides the ability to capture and identify kinases from whole cell lysates on a kinome scale. Binding of kinases is dependent on the functional expression and activity of the kinase and affinity for the different immobilized inhibitors. To determine the inhibitor bead selective distribution of bound kinases, we assayed kinase capture by six different inhibitors individually covalently coupled to Sepharose beads [7,9]: CTx-0294885, VI-16832, PP58, Purvalanol B, and two custom synthesized molecules, UNC-8088A and UNC-2147A. Four cell lines representative of breast cancer subtypes: HCC1806 (basal-like), SUM159 (claudin-low), MCF7 (luminal), and SKBR-3 (HER2-enriched) were used for analysis of kinase capture by each bead (Figure 1A). Of these, CTx-0294885 (CTx) and VI-16832 (VI) captured the most total kinases (265 and 254, respectively) and the most unique kinases (32 and 29, respectively). The other four beads bound a lesser number of kinases (PP58, 194 kinases; Purvalanol B, 164; UNC-8088A, 162; UNC-2147A, 130, Figure 1B, Table S1). Although UNC-8088A binds the fewest unique kinases (only five), these include the atypical bromodomain and extraterminal (BET) domain-containing family of chromatin readers BRD2, -3, and -4 [10]. Hierarchical clustering of identified kinase peptides shows each bead binds a unique set of kinases and UNC-2147A displays the most distinct binding profile selectively enriching the AGC kinases (Figure 1C).

**Fig. 1.**
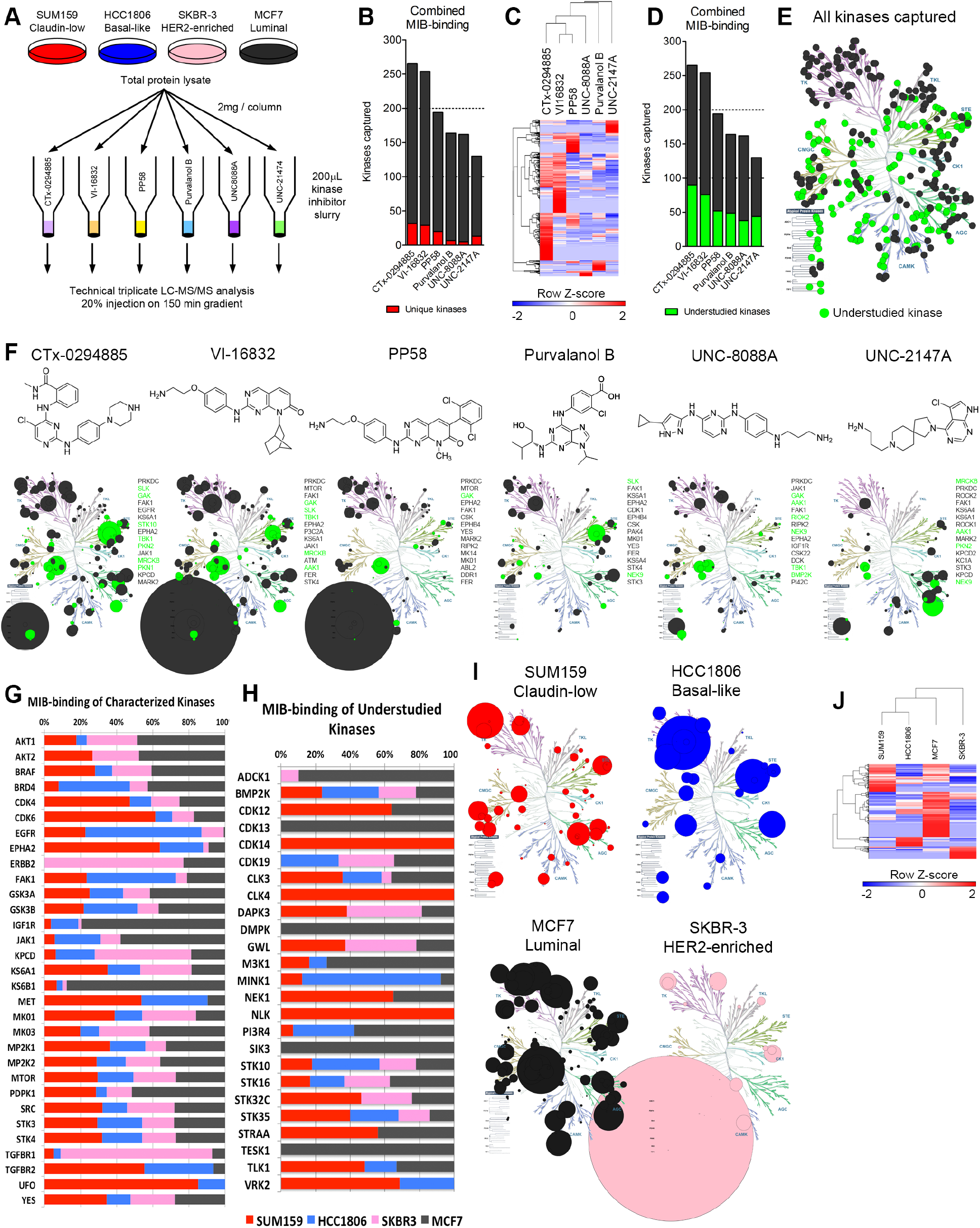
Assessment of multiplexed kinase inhibitor beads (MIBs) for kinase capture across breast cancer subtypes. (A) Experimental design to assess performance of six kinase inhibitor beads. (B) Combined data from all four cell lines assayed shows CTx-0294885 binds the most number of kinases. Number of kinases captured uniquely by each bead is shown in red. (C) Euclidean hierarchical clustering kinase peptides bound by the six beads shows each bead enriches for a distinct set of kinases. UNC-2147A displays the most unique binding profile. (D) A large proportion of kinases captured by MIBs (23-24%) are understudied or poorly characterized (green). (E) 381 kinases were identified across all four cell lines, including 346 protein kinases and 35 metabolic kinases. Of these protein kinases, 142 are understudied (green). (F) Chemical structures and kinase-binding of each inhibitor bead across the kinome. Circle size is proportional to the number of unique peptides identified per kinase. PRKDC (DNA-PK) is over-represented in VI-16832 and PP58 (large circle under Atypical protein kinases). Most beads capture kinases across families but UNC-2147 preferentially enriches for AGC family kinases. Shown to the right of each kinome tree are the 15 most-highly captured kinases for each bead. Green circles and text signify understudied kinases. (G) Comparison of relative binding of characterized kinases across breast cancer cell lines/subtypes. (H) Comparison of relative binding of understudied kinases across breast cancer cell lines/subtypes. (I) Each cell line representing the different breast cancer subtypes displays a unique kinome profile. Only kinases with the greatest number of peptides identified in particular cell line are shown. Circle size is proportional to the number of peptides identified. (J) Hierarchical clustering of peptides identified for each kinase (rows) across the cell lines (columns) cluster triple-negative cell lines (SUM159, HCC1806) together and indicates HER2-positive SKBR3 cells have the most distinct kinome profile.

Understudied kinases [2] make up approximately 40% of the overall expressed kinome and are similarly represented through MIB-binding, with 23-34% of all kinases captured for any individual bead (Figure 1D; Table S2). Characteristics of understudied kinases includes: i) integration of the protein kinase in signaling networks is poorly defined, ii) function and/or regulation is poorly defined, iii) activation loop phospho-antibodies and/or IHC grade antibodies may not exist, iv) lack of selective chemical tools for use in characterization of function (e.g., small molecule inhibitors), v) RNAi and CRISPR/Cas9 for knockout/altered expression and cDNAs for overexpression may be primary tools, vi) kinase knockout or altered expression may not provide readily assayable phenotypes (e.g., growth, migration, apoptosis or in vivo function in mouse organ physiology).

Across all MIB/MS runs, 381 kinases in total were identified. Of these, 35 are metabolic and lipid kinases, 346 are protein kinases of which 142 can be considered understudied (41% of protein kinases identified) (Figure 1E). The overall distribution of kinases bound indicates CTx and VI are clearly pan-kinase inhibitors (Figure 1F, circle size proportional to number of unique peptides identified per kinase). Purvalanol B also binds kinases across families but to a lesser extent. PP58 has some preference for tyrosine kinases (TKs), and UNC-8088A has preference for TKs, CMGC, and atypical kinases over other families. UNC-2147A, designed for interaction with the binding pocket of AKT, has a strong affinity for AGC kinases lacking in most of the other five kinase inhibitors. CTx, VI, and PP58 have a strong affinity for the PRKDC (DNA-PK) not seen with the other three inhibitors. All inhibitor beads display high affinity for many understudied kinases (green circles and text). The most-highly captured understudied kinases across the four cell lines were GAK, SLK, MRCKB, AAK1, TBK1, and NEK9.

Kinases known to be oncogenic drivers in general and/or nodal signaling kinases display anticipated MIB-binding profiles across the different breast cancer subtype cell lines (Figure 1G). For example, SKBR3 (luminal HER2+) and MCF7 (luminal) cells have abundant AKT1/2 MIB-binding. Other well-characterized kinases are highly represented in a specific cell line, such as EGFR and FAK1 in HCC1806, EPHA2 and UFO (AXL) in SUM159, IGF1R and KS6B1 (p70 S6K) in MCF7, and HER2/ERBB2 and TGFBR1 in SKBR-3 cells. Several understudied kinases also show high selectivity in functional MIB-binding including CDK13, DMPK, SIK3 and TESK1 in MCF7 and CLK4, CDK14 and NLK in SUM159 cells (Figure 1H). Figure 1I and Table S1 show kinases whose MIB-binding is greatest in each of the four cell lines, proportional to the number of unique peptides identified. Unsupervised hierarchical clustering illustrates the differences in MIB-binding throughout the kinome for each cell line (Figure 1J). These findings indicate the four cell lines display a unique MIB/MS binding profile for both well-characterized and understudied kinases.

### Integrating understudied and well-characterized kinases by kinome proteomic profiling defines breast cancer subtypes

We characterized the baseline kinome of 15 breast cancer cell lines representing the four major breast cancer subtypes previously defined by gene expression profiles [3]. Cell lysates were passed over an affinity column composed of the six kinase inhibitor beads and processed for LC-MS/MS (Figure 2A). Using label-free peptide quantitation measurements a total of 360 kinases were identified having at least 3 unique peptides (Supplemental Data file S1). Claudin-low and basal-like cells (TNBC) are readily distinguished from HER2-enriched/luminal cells by MIB profiling of the cellular kinomes shown by consensus clustering (Figure 2B). The basal-like HER2-amplified cell line HCC1954 clusters with basal-like lines through similarity of kinome profiles and is thus separated from the luminal HER2+ lines. Interestingly, the SKBR-3 HER2-enriched cell line shows an intermediate clustering between HCC1954 and other HER2+/luminal cell lines, and a previous report demonstrated SKBR-3 patterns as basal-like in functional RNAi screens [11]. Hierarchical clustering of kinases further separated cell lines with the claudin-low phenotype, SUM159, MDA-MB-231 and MDA-MB-468 (basal-like), showing the greatest difference from other cell lines (Figure 2C). SUM229 cells have two subpopulations, a basal-like EpCAM positive/E-cadherin positive (SUM229pos) and a claudin-low EpCAM negative/E-cadherin negative population (SUM229neg). The two populations are genomically similar by exome sequencing, but differ epigenetically [12] and cluster together based on their kinome MIB-binding profile (Figure 2C).

**Fig. 2.**
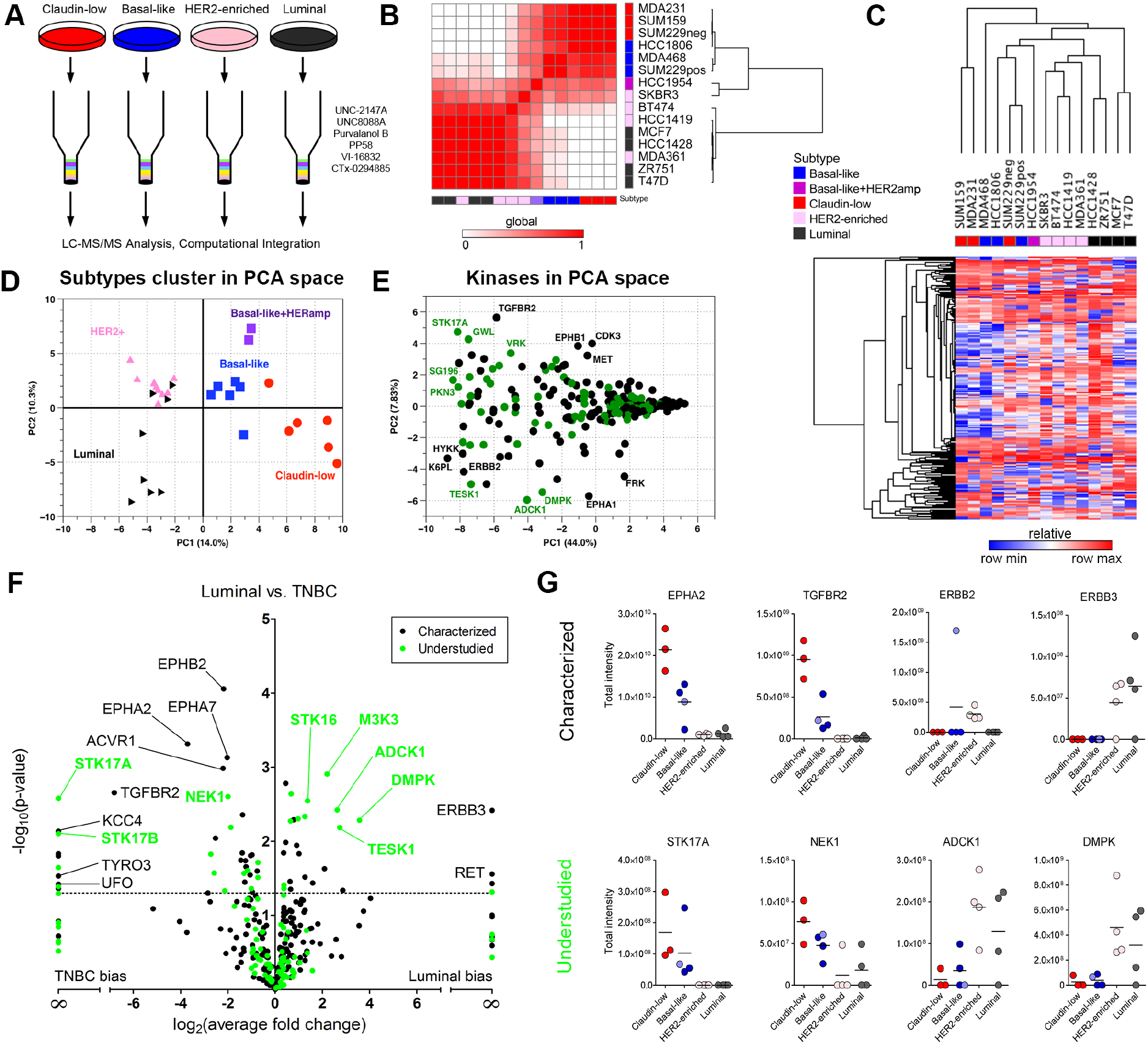
MIB/MS kinome profiling assigns breast cancer cell lines to functional subtypes. (A) Individual samples run across 6-bead composition with LC-MS/MS analysis. (B) Heat map of correlation between MIB/MS samples for cell lines analyzed. Color bars indicated the subtype of each cell line (blue: basal-like, red: claudin-low, pink: HER2-enriched, black: luminal, purple: basal-like/HER2amp). White in the heat map indicates a low correlation between samples, while red shows higher correlation. Rows and columns are hierarchically clustered. (C) Heat map of MIB/MS average for each of the 15 cell lines analyzed. Rows are kinases; columns are MIB/MS cell line averages. Color bar for columns indicates the subtype associated to each cell line. Each column is an average of 2 or 3 MIB/MS samples, depending on the cell line. Colors in the heat map are relative by row minimum (blue) and maximum (red). A total of 254 kinases passed filtering (see Methods). Rows and columns are hierarchically clustered using Euclidean distance and average linkage. (D) Principal Component Analysis (PCA) on the entire MIB/MS data set. PC1 and PC2 account for 14.0% and 10.3% of the variance in the data set, respectively. A total of 32 samples across the fours subtypes are represented by their subtype (red circle: claudin-low, blue square: basal-like, pink upward triangle: HER2-enriched, black right-pointing triangle: luminal, purple square: basal-like/HER2amp). (E) PCA on the MIB/MS data set to show highly variable kinases across the samples. Characterized and understudied kinases are shown in black and green, respectively. PC1 and PC2 account for 44.0% and 7.83% of the variance in the data set, respectively. (F) Volcano plot showing characterized (black) and understudied (green) kinases that are significantly (p<0.05) different between the Luminal/HER2-enriched and TNBC (basal-like/claudin-low) cell line samples in the MIB/MS data set. (G) Profiles of selected characterized (top row) and understudied (bottom row) kinases across breast cancer subtypes.

Principal components analysis (PCA) of baseline MIB-binding kinase profiles revealed significant differences between subtypes within the first principal component, clearly separating triple-negative from HER2-enriched and luminal cell lines (Figure 2D). Further separation of the triple-negative group into claudin-low and basal subtypes is also readily achieved. Appreciable separation of HER2-enriched cell lines from luminal cell lines is observed in the second principal component, as is that of the basal-like/HER2-amplified cell lines from the basal-like and claudin-low lines. A loadings plot, which defines relationships between MIB-binding for each kinase, highlights those kinases with significant variation within subtypes, with numerous understudied kinases being apparent (Figure 2E). Examples of understudied kinases with differences in MIB-binding among cell lines include ADCK1, PKN3, STK17A and TESK1. Similarly, well-characterized kinases known for their relevance in breast cancer are observed, such as ERBB2, EPHA1, MET and TGFBR2.

Supervised differential expression analysis of MIB-captured kinases from claudin-low/basal-like (TNBC) versus HER2/luminal cell lines defined several statistically significant differences (Figure 2F). Multiple Ephrin receptors (EPHA2/A7/B2) and members of the TGF-beta superfamily (TGFBR2, ACVR1) are among the kinases most associated with TNBC while ERBB3 and RET are over-represented in HER2+ and luminal cell lines. Many understudied kinases display higher MIB-binding in HER2+/luminal cells, including DMPK, ADCK1, and TESK1. Individual plots for selected kinases are shown in Figure 2G, showing distinctive patterns of MIB-binding across subtypes.

### Kinase MIB-binding activity is independent from mRNA expression level

Our results clearly demonstrate that kinase MIB-binding displays strong variation across breast cancer subtypes. Global gene expression measurements have similarly shown subtype-specific dynamics, with expression of selected gene sets being utilized in subtype determination and diagnosis [13–15]. We compared baseline RNA-seq measurements with corresponding MIB-capture of protein kinases to assess the relationship between transcript abundance and functional kinome behavior. Similarity of kinase profiles for a given measurement modality was highly similar, with unsupervised hierarchical clustering grouping RNA-seq profiles separately from those derived from MIB-binding (Figure 3A). Furthermore, similarity within a modality was very high, such that breast cancer subtypes were largely clustered correctly, especially when looking at MIB-binding profiles that clearly grouped along basal, HER2+ and triple negative subtypes.

**Fig. 3.**
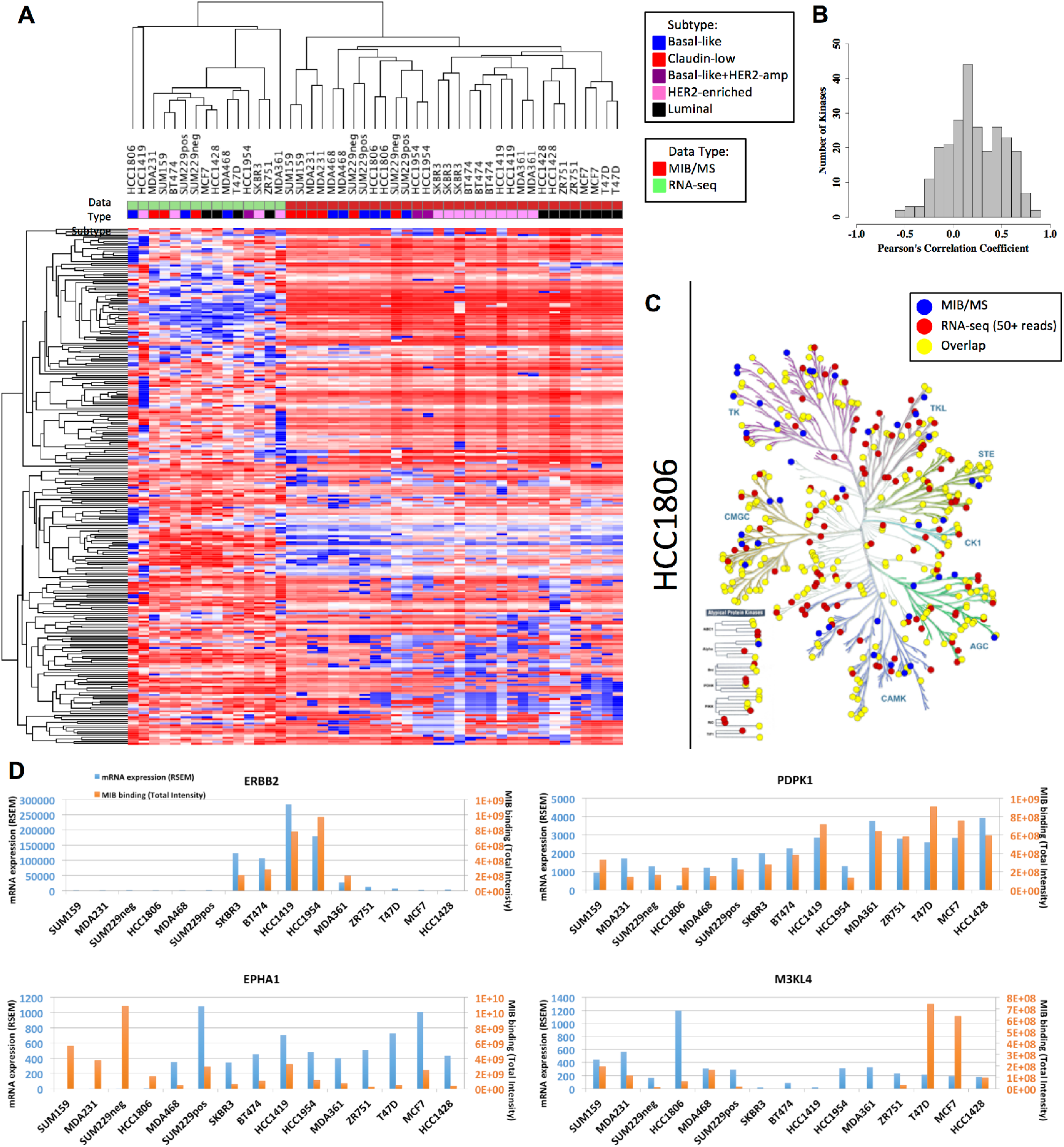
Overall MIB-binding and mRNA expression levels are not correlated. (A) Heatmap of autoscaled mRNA expression and MIB-binding for all of the samples analyzed. Rows and columns are hierarchically clustered. Rows are the 254 kinases. The “Data Type” track along the columns indicates which type of data the sample is from (green: RNA-seq or red: MIB/MS). The “Subtype” track along the top of the heatmap indicates which subtype the cell line is classified as (blue: Basal-like, red: Claudin-low, purple: Basal-like+HER2-amp, pink: HER2-enriched, and black: Luminal). (B) Frequency distribution of Pearson’s correlation coefficient across all cell lines in MIB/MS and RNA-seq for each of the 254 kinases. (C) KinomeTree for MIB-binding (blue dots), RNA expression (red dots), and the overlap between the two data sets (yellow dots) for the HCC1806 (basal-like) cell line. (D) Representative raw profiles of ERBB2, PDPK1, EPHA1, and M3KL4 in MIB/MS intensity (orange bars) and mRNA RSEM counts (blue bars), showing both highly correlated and poorly correlated behavior between the two data sets.

While within-group correlations were high, normalized RNA-seq was found to have a very poor correspondence to MIB-binding using label-free quantification of kinase peptide abundance. Quantitative comparison for each subtype achieved correlation coefficients of no more than 0.25, implying that less than 7% of the observed variation between MIB-binding and RNA abundance in breast cancer subtypes is explained through this relationship. The distribution of the Pearson correlation coefficients of all kinases across all 15 cell lines similarly shows a low correspondence between MIB-binding and RNA-seq (Figure 3B), with the mean correlation being 0.2. These results are consistent with other work that found the average correlation between gene expression and protein abundance in TCGA colorectal cancer samples to be approximately 0.47, with a lesser correlation of 0.23 when comparing gene expression and protein variation [16]. A more recent comprehensive analysis of several data sets has further shown that mRNA levels are not predictive of protein levels for a given gene [17]. The low correlation between RNA-seq and MIB-binding suggests that the use of MIB/MS provides a picture of kinome behavior complementary to that provided through RNA expression measures. In particular, these results support the potentially significant role of post-translational and transcriptional regulation in kinome dynamics [17], [18], [19].

While there is significant overlap, there are a number of kinases that are only observed with one of the applied methods, MIB/MS or RNA-seq (Figure 3C). This discrepancy is partly due to the 50+ RSEM read threshold used here as a positive identification in RNA-seq, potentially missing very lowly expressed kinases. Similarly, kinases not observed with MIB/MS but identified in RNA-seq may be missed due to being in an inactive, nonfunctional state and/or failure of chosen inhibitors to bind these kinases with adequate affinity. Pseudokinases that do not bind ATP will generally not be captured by MIBs.

While the degree of correlation between RNA-seq and MIB-binding can vary significantly for a given kinase, we do observe a broad range of behaviors across cell lines, with representative high and low correlation profiles shown in Figure 3D. ERBB2 is strongly expressed in HER2+ cell lines and MIB-binding is correspondingly strong in these samples, consistent with the importance of transcriptional regulation in this kinase’s functional output. PDPK1 similarly shows high correlation of expression and MIB-binding, though across all cell lines, while conversely EPHA1 and M3KL4 (MAP3K21) show very poor correlations.

Such proteomic behavioral properties cannot be detected by RNA-seq alone. Together, MIB/MS coupled with RNA-seq provides an integrated perspective, providing a post-transcriptional measure of kinase protein levels. While the functional consequences of post-transcriptional regulation in relation to kinase networks and signaling is not understood, the observed differential relationship of transcript versus protein expression for a subset of poorly correlated kinases suggests an unknown control mechanism possibly involving differential covalent regulatory modifications and/or protein stability. Importantly, MIB/MS measurements avoid the use of recombinant kinases, often used to profile on-target/off-target inhibitor profiles that are not representative of endogenous kinase complexes [20,21]. Using MIB capture and MS quantification of endogenous kinases in cell lysates that have associated regulatory subunits and post-translational modifications thus provides a functional measure of cellular kinase protein expression.

### Kinome profiles accurately define tumor biopsies

Cumulatively, our data show that measurement of kinases by MIBs capture allows integration of a significant fraction of the expressed kinome, defining a taxonomy of breast cancer determined by the functional behavior of protein kinases. This kinome taxonomy is used below to define a subset of understudied and well-characterized kinases that are capable of distinguishing between breast cancer tumor subtypes.

Given both the variation in kinase MIB-binding profiles observed across subtypes, as well as their differing information content when compared to RNA expression measurements, we sought to better understand which kinases were key nodes in the subtype-selective baseline breast cancer kinome. To address this question, we investigated the MIB-binding behavior of kinases across all four subtypes. We considered three major classes of kinases: 1) those that show variation in MIB-binding across all subtypes, 2) those that exhibit more limited subtype-specific behaviors, and 3) kinases that have nominal distinguishing behavior. Standard application of PCA identifies those kinases displaying the greatest variation across all samples (“pan-subtype kinases”; Figure 2E) and thus we used a feature selection approach based on the Bhattacharyya distance [22] to determine subtype-specific kinases that are highly distinguishing/informative for a single cancer subtype (see Methods). Integrated with PCA-identified kinases (“pan-subtype”), this combined set of the 50 most informative kinases is shown in Figure 4A, with column ordering based on similarity of the kinome profile and recapitulating similarity between claudin-low and basal subtypes as well as HER2-enriched and luminal. The HER2+ cell line that profiles as basal-like (HCC1954, in purple) is displayed separately. Recognized cancer-related kinases are again observed in this set, including ERBB2, FGFR2, PTK6, RAF1 and RON (MST1R) as well as 22 understudied kinases. Using the pan-and subtype-specific kinases in Figure 4A, we assessed their effectiveness in subtype identification using a support vector machine (SVM) classifier along with leave-one-out cross-validation. We found that the ability to classify a cell line’s subtype from measurement of these kinases was highly accurate in both precision and specificity, especially for claudin-low and basal subtypes (Table S3).

**Fig. 4.**
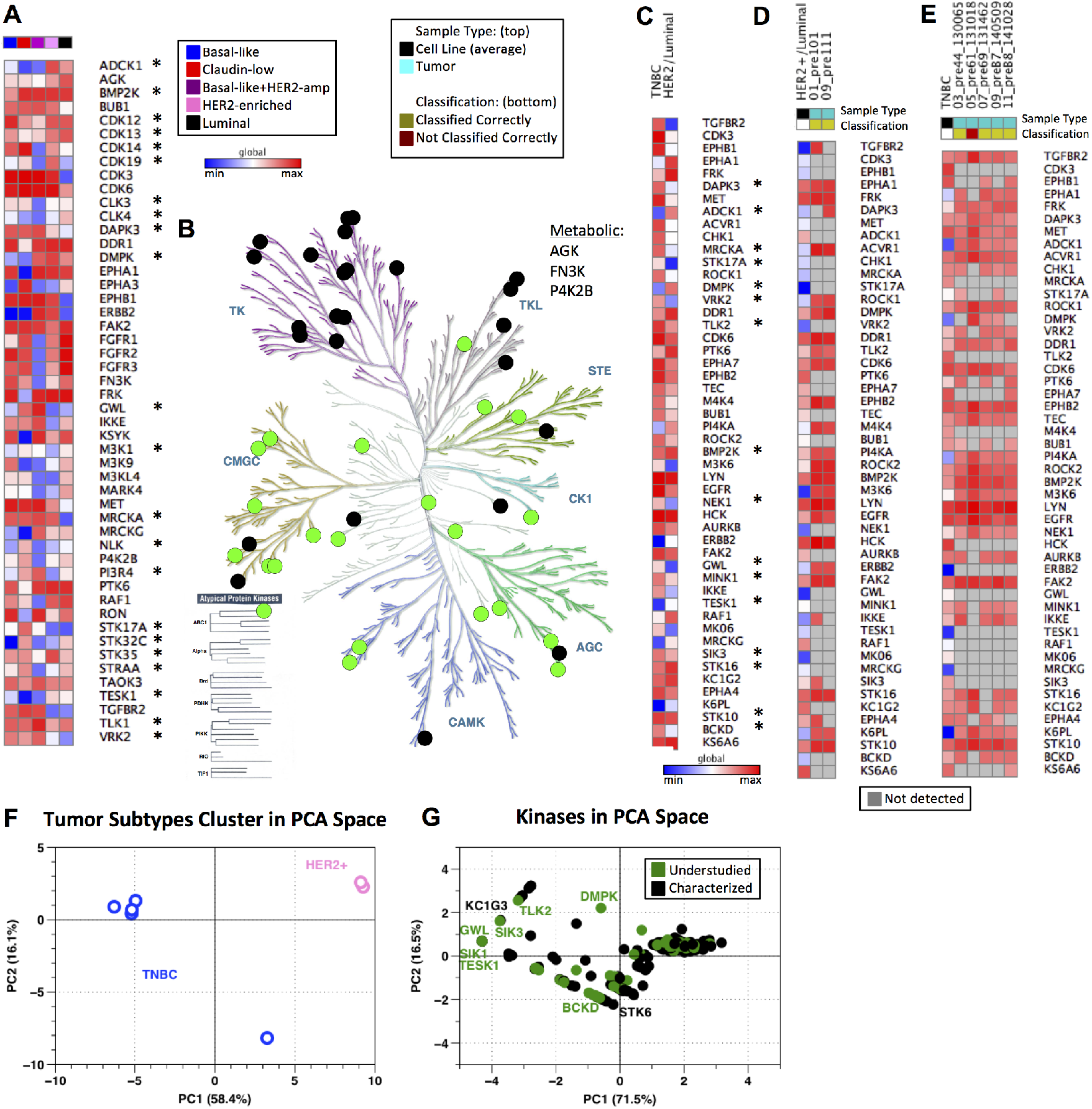
Baseline kinome of cell lines and tumors across breast cancer subtypes. (A) Compilation of subtype specific and pan-subtype kinases chosen from feature selection and PCA, respectively. All data is log2 normalized and autoscaled by sample, with heat map colors indicating low (blue) to high (red) MIB-binding. Column color bar indicates subtype (red: claudin-low, blue: basal-like, pink: HER2-enriched, black: luminal, purple: basal-like/HER2amp; Understudied kinases are denoted by *). Global maximum and minimum color assignment. (B) KinomeTree with the 50 distinguishing features from (A) are denoted. Black circles denote characterized kinases, while green circles represent understudied kinases. (C) Kinases chosen from feature selection when comparing Luminal/HER2-enriched cell line samples against basal-like/claudin-low (TNBC) cell line samples. Kinases are ordered from top to bottom in the same ordering as from the feature selection (most heavily weighted kinases are at the top of the heat map). All data is log2 normalized and autoscaled by sample, with heat map colors consistent with those in (A) (Understudied kinases are denoted by *). Global maximum and minimum color assignment. (D) Heat map of Luminal/HER2-enriched cell line average (HER2+/Luminal column; black in “Sample Type” column color bar) across the kinases shown in (C) with two tumor samples (teal in “Sample Type” column color bar). Data is log2 normalized and autoscaled by samples, as previously noted. Yellow in the “Classification” column bar shows which samples are classified correctly as Luminal/HER2-enriched by the SVM using the kinases from (C). Blue in the heat map indicates a low MIB-binding, red indicates high MIB-binding, and grey (in the tumor samples only) indicates that a kinase was not detected by MIBs in the tumor sample. Global maximum and minimum color assignment. (E) Heat map of TNBC cell line average (TNBC column; black in “Sample Type” column color bar) across the kinases shown in (C) with five tumor samples (teal in “Sample Type” column color bar). Data is log2 normalized and autoscaled by samples, as previously noted. Yellow in the “Classification” column color bar shows which samples are classified correctly as TNBC by the SVM using the kinases from (C). Dark red in “Classification” indicates that the tumor sample was incorrectly classified (not classified as TNBC) by the SVM using the kinases identified in (C). Color scheme in the heatmap is consistent with that described in (D). Global maximum and minimum color assignment. (F) PCA scores plot of tumor samples with PC1 and PC2 accounting for 58.4% and 16.1% of variance, respectively. TNBC tumors are blue and HER2-enriched tumors are pink. (G) PCA loadings plot of tumor samples with PC1 and PC2 accounting for 71.5% and 16.5% of variance, respectively. Black points are characterized kinases and green points denote understudied kinases.

The kinases shown in Figure 4A have the greatest variation within and across subtypes and are representative of each of the major subfamilies of kinases in addition to three metabolic kinases captured by MIBs (Figure 4B). Under the assumption that TNBC (represented by the basal and claudin-low cell lines) and HER2/luminal breast cancer are separate diseases, we again used unsupervised feature selection of MIB/MS data to identify kinases that distinguish TNBC (basal/claudin-low) from HER2/luminal breast cancer [22]. As shown in Figure 4C, obvious differences in the kinome profiles of TNBC and HER2/luminal are observed, demonstrating the unique functional phenotypic features of the kinome in the two different breast cancers. Sixteen understudied kinases showed strong variance between TNBC and HER2/luminal breast cancer, with higher-ranked understudied kinases being DAPK3, ADCK1, MRCKA (CDC42BPA), STK17A, DMPK and VRK2.

Using the kinases chosen from feature selection in Figure 4C, we evaluated the ability to use MIB-binding profiles to define subtypes of human HER2+ needle biopsies and TNBC breast tumors (Figure 4D & E). Diagnostic needle biopsies of 2 patient tumors (2 HER2+) having ~1 mg of total protein were processed using MIB enrichment. With just 1 mg of tumor lysate protein, the total number of kinases purified from each biopsy ranged from ~200 to 275. Utilizing only the kinases identified from cell lines as the identifying features (Figure 4C) within a SVM classifier, it was possible to clearly identify HER2+ and TNBC primary patient tumors. As with cell lines, application of PCA to MIB-binding profiles showed a clear separation between TNBC tumors and HER2+ tumors (Figure 4F). Kinases driving the variation within the data included ERBB2 as expected and several understudied kinases including TESK1 and DMPK (Figure 4G). Thus, the MIB-binding activity of no more than 50 kinases is sufficient to discriminate the functional phenotypic nature of the kinome in breast cancer.

### A functional interaction network of MIB-binding kinases

To establish a basic picture of the architecture of the human kinome using MIB-bound kinases, we compiled protein interaction and phosphorylation data from multiple data sources and established a functional interaction network among 246 of the 254 kinases commonly identified in the panel of 15 breast cancer cell lines (see Methods). Spectral clustering of this network further enabled the identification of 16 subnetworks, with many showing functional enrichment of one or more Gene Ontology functional categories (Figure 5A; GO term enrichment for subnetworks is provided in Supplemental Data File S4). Of note is that understudied kinases (green nodes) are widely distributed across all the major subnetworks, demonstrating that these poorly characterized kinases are integrated into subnetworks along with well-characterized kinases. The 50 distinguishing kinases identified for cell lines in Figure 4A (triangle nodes) were also distributed throughout the network and associated subnetworks. The breadth of subnetwork coverage by these kinases suggests that their predictive value in our subtype classification comes from their distribution across many subnetworks, providing an overall estimate of the state of many functional processes simultaneously.

**Fig. 5.**
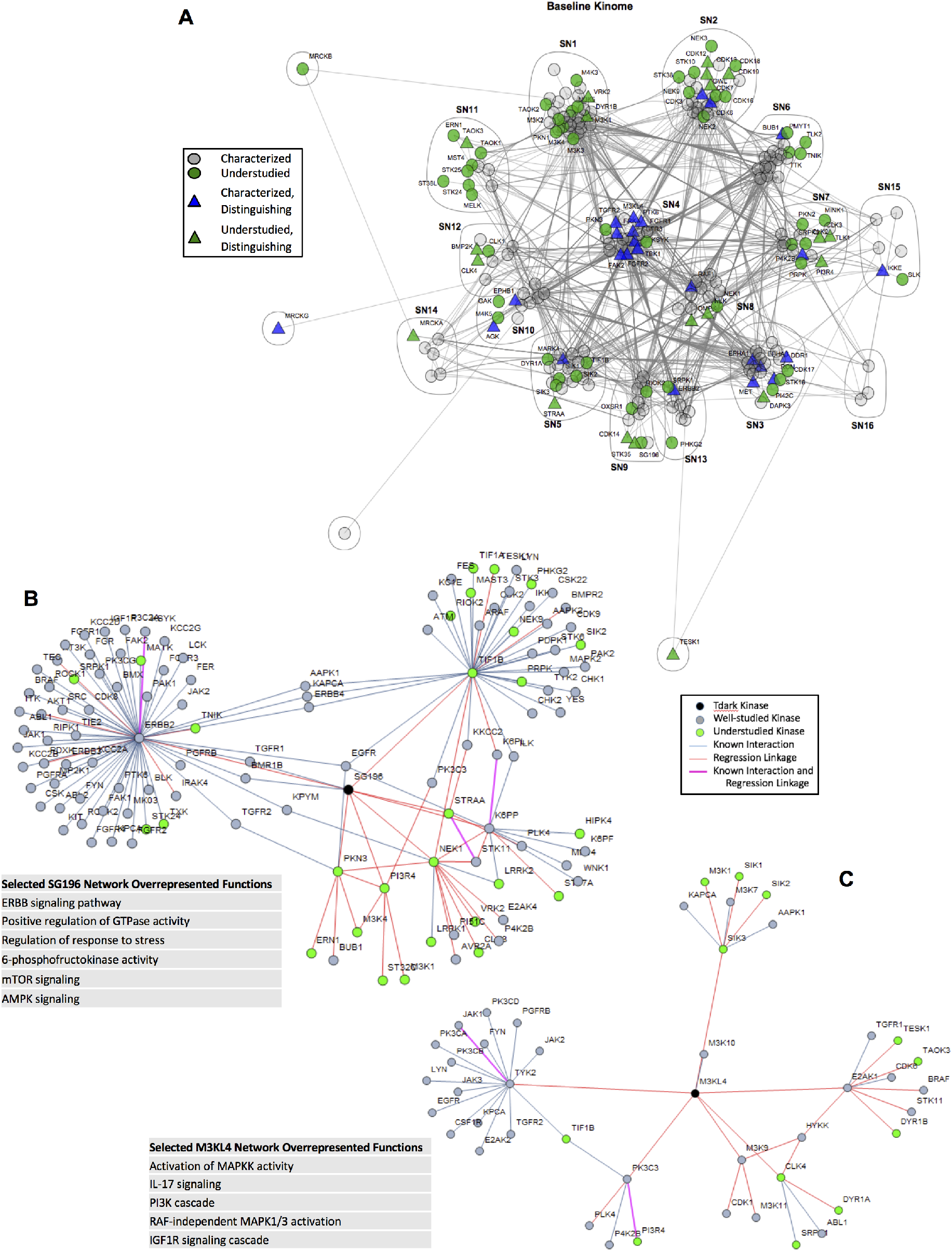
Subnetworks in the functional kinome. (A) Compiled and spectral clustered protein-protein interaction network from public data sources of the 254 kinases analyzed in the MIB/MS cell line data set. Green nodes represent understudied kinases, while grey and blue nodes represent well-characterized kinases. Triangles are kinases that are also in the distinguishing features found in Figure 4A. (B & C) Local functional network for SG196 (B) and M3KL4 (C) as defined through Lasso regression of MIB/MS data with sample enriched annotations from GO, Kegg and Reactome pathways.

Of the kinases bound by MIBs, we identified four Tdark kinases within our data, including ADCK1, CSK23, SG196 and M3KL4 (MAP3K21). As defined in the NIH Illuminating the Druggable Genome (IDG) program, Tdark kinases are very poorly characterized in terms of publications, small molecule inhibitors and gene references to functionality [1]. To provide an example of how MIB/MS can be used to help provide putative functional roles for such poorly characterized kinases, we utilized Lasso regression to identify potential functional linkages for these kinases as well as all understudied kinases, including those identified as Tdark and Tbio by the IDG program. As described in greater detail in Methods, known physical interactions were first identified from multiple data sources for all kinases. These were then integrated with potential functional linkages between kinases identified through Lasso regression across MIB-binding profiles. Together, these data established functional subnetworks centered on each individual understudied kinase. In total, 89 understudied kinases were annotated with such functional subnetworks, of which 18 were defined as Tbio and 4 as Tdark kinases. Functional annotation of these groups was then performed to identify enriched functional GO categories or signaling pathways, with results for all 89 kinases provided in Supplemental Data File S5.

An example of this analysis as applied to the Tdark kinase SG196 (Sugen Kinase 196 or protein-O-mannose kinase) is shown in Figure 5B. SG196 was found to have two known physical interactions in addition to eight regression linkages that reveal over-representations of the ERBB signaling pathway, MAPK cascade, and positive regulation of GTPase activity as well as several other statistically enriched biological functions as identified through GO and pathway analysis of the entire functional subnetwork identified in Figure 5B. Similarly, M3KL4 (MAP3K21) has only a single known physical interaction but by utilizing the correlated behavior of the regression kinases the functional network has overrepresentation of MAPKK activity, regulation of immune system, and response to stress (Figure 5C). The networks and functional annotations for the other Tdark kinases, CSK23 and ADCK1 are provided in Supplemental Data File S5.

### Kinome MIB-binding profile and response to drug perturbation

To assess how the baseline kinome and associated understudied kinases change in their functional MIB-binding profile in response to targeted drug perturbation, we exposed four cell lines to three subtype-relevant kinase inhibitors: SUM159 and HCC1806 with trametinib (a MEK1/2 inhibitor); SKBR3 with lapatinib (a HER2/EGFR inhibitor); and MCF7 with buparlisib (a PI3K inhibitor). Each inhibitor strongly suppressed growth of the selected subtype specific cell line (Figure 6A). We have previously shown that the kinome is dynamic and rapidly adapts to targeted perturbation by kinase inhibitors [6,7]. This adaptive response is readily observed by changes in the MIB-binding profiles for each drug treatment (Figure 6B, Supplemental Data File S3), with SUM159 cells showing the strongest dynamic response to drug perturbation relative to the other cell lines, but each line clearly shows an adaptive response of the kinome measured by MIB-binding profiles. Figure 6C shows the kinases that are unique to each subtype defined in Figure 2C (Table S4).

**Fig. 6.**
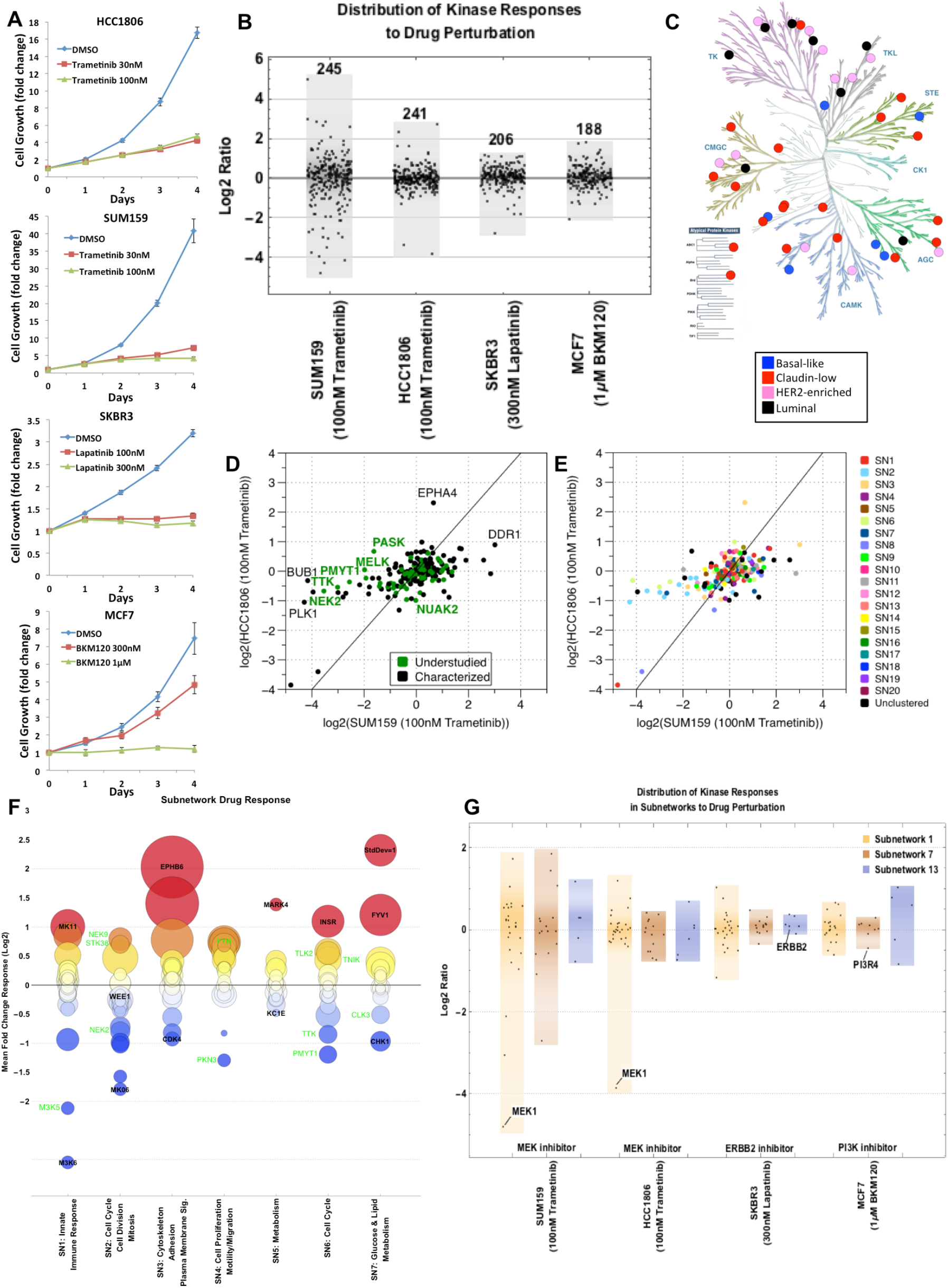
Kinome drug response overall and by subnetwork. (A) Growth curves for HCC1806 + Trametinib, SUM159 + Trametinib, SKBR3 + Lapatinib, and MCF7 + BKM120. All curves were done at two different doses. (B) Response of kinome in representative cell lines across four subtypes of breast cancer (claudin-low: SUM159, basal-like: HCC1806, HER2-enriched: SKBR3, luminal: MCF7) when treated with the indicated kinase inhibitor. Distribution of the kinome response on the log2-scale is shown for each cell line/subtype; each point represents a kinase. (C) KinomeTree showing the kinases that are uniquely captured in each of the subtypes in the baseline data set. Blue circles denote kinases bound to the MIBs only in basal-like samples. Similarly, red circles represent claudin-low, pink circles represent HER2-enriched, and black circles represent luminal uniquely bound kinases. (D) Scatter plot of the response of the basal-like vs. claudin-low cell lines to treatment with 100nM Trametinib. All values are fold change to untreated cells and log2-transformed. Kinase points are colored black for characterized and green for understudied. (E) Scatter plot of the response of the basal-like vs. claudin-low cell lines to treatment with 100nM Trametinib (same as in (D)). Kinases are colored by subnetwork assigned to each kinase from Figure 5. (F) Subnetwork response to drug perturbation showing mean fold change across the four representative cell lines (SUM159, HCC1806, SKBR3, and MCF7) for the top 7 subnetworks identified from Figure 5. Characterized and understudied kinases in each subnetwork are labeled in black and green, respectively. The color of each circle indicates the mean fold change (red=high/above 0, blue=low/below 0), while the area of the circle denotes the standard deviation of the fold changes across the representative cell lines. (G) Distribution of the kinome response in the three subnetworks (SN1, SN7, and SN13 on the log2-scale is show for each cell line/subtype.

Scatter plots of the SUM159 and HCC1806 dataset defines specific kinases and kinome subnetworks that drive the adaptive response to MEK1/2 inhibition that are represented by both understudied and well-characterized kinases (Figure 6D). Understudied (i.e. NEK2 and PASK) and well-characterized (i.e. DDR1 and EPHA4) kinases respond differently in the two subtypes (basal-like and claudin-low) when they are treated with the same kinase inhibitor (trametinib). The kinases in the subnetworks defined in Figure 5 also respond uniquely in the basal-like and claudin-low subtypes when treated with trametinib (Figure 6E). The adaptive kinome response measured by dynamic changes in MIB-binding profiles is more clearly seen when specific subnetworks are analyzed (Figure 6F). The global response of the seven largest subnetworks to these drug perturbations is shown with their functional annotation as estimated from Gene Ontology terms and KEGG pathway enrichment shown on the x-axis and MIB-binding response reported on the y-axis as a mean across all cell lines and drugs. Subnetworks have heterogeneous responses, with some subnetworks being fairly coordinated in response and others having kinase members acting in a more strongly divergent manner. For instance, subnetwork 3 (SN3) is enriched with many kinases relevant to cytoskeleton, adhesion and motility and has many of its members strongly up-regulated in response to drug perturbation. In comparison, SN2, involved in cell cycle and cell division, contains both strongly up-regulated and down-regulated kinases, with the largest responses being loss of MIB-binding, consistent with the inhibition of cell growth. Understudied kinases (green labels in Figure 6F) often display large responses to drug treatment within a given subnetwork, demonstrating these kinases actively contribute to adaptive kinome reprogramming in response to targeted kinase inhibition. Similarly, a more detailed look at targeted inhibition of specific subnetworks for each of the cell lines shows the dynamic response of the kinome to be highly dependent on the drug, subtype and subnetwork context (Figure 6G, Table S5). More broadly, the response of kinases in subnetworks is consistent with a unique functional regulation of the kinome in cancer subtypes and in response to different perturbations.

## Discussion

While the creation of molecular taxonomies has established the existence of subtypes in many tissue-specific cancers, how these taxonomies can be leveraged to characterize phenotype or to guide the development of targeted therapeutics remains unclear. A complication for improving therapeutic intervention with targeted kinase inhibitors in cancer is the extensive number of understudied kinases, whose poor characterization presents significant challenges to understanding their role in emergent processes such as adaptive bypass reprogramming and resistance to kinase inhibitors. Despite such challenges, understudied kinases do have potential as novel drug targets once their functional integration into signaling networks is more clearly determined. Methods generally have been lacking to capture kinases, both well-characterized and understudied, to define the functional kinome en masse. Characterization of kinase MIB-binding in tumor cell lysates has proven to be a powerful technique for characterizing functional architectures of the kinome that provides the capability to identify prognostic signatures and differential response to perturbations such as targeted kinase inhibition, as well as better establishing the integration and function of understudied kinases. This is clearly seen in the 50 kinase profile distinguishing TNBC from HER2+/luminal breast cancer with many of the 50 kinases representing understudied kinases.

The highest weighted understudied kinases distinguishing TNBC from HER2+/luminal breast cancer include ADCK1 (AarF Domain Containing Kinase whose function is unclear), DAPK3 (Death-associated protein kinase thought to be involved in apoptosis), DMPK (Dystrophia myotonica protein kinase whose function is not well-defined), MRCKA (Myotonic dystrophy kinase-related CDC42 binding protein kinase alpha that may signal CDC42 control of the actin cytoskeleton and is related to DMPK), STK17A (Serine/threonine kinase 17A has apoptosis-inducing activity and is a member of the DAP kinase-related family), TLK2 (Tousled-like kinase 2 is involved in chromatin assembly and possibly DNA repair) and VRK2 (Vaccinia-related kinase 2 that is believed to regulate apoptosis and cell growth). Screening of the cBioPortal for Cancer Genomics (http://www.cbioportal.org/public-portal/) indicates amplification of MRCKA (CDC42BPA) in 13-25% of invasive breast cancer while TLK2 is amplified in 10% of invasive breast cancer and 25% of adenoid cystic breast cancer. ADCK1, DAPK3, DMPK STK17A and VRK2 were found to be similarly amplified in other cancers including prostate adenocarcinoma, uterine carcinosarcoma and pancreatic adenocarcinoma. Prominent MIB-binding signatures combined with potential increased expression in tumors suggests these understudied kinases have important functions for the tumor cell phenotype that have not been characterized to date.

The dynamic nature of the kinome is clearly captured in the kinase MIB-binding profiles characterizing baseline versus post-drug treated cells. This adaptive reprogramming of the kinome is involved in the epigenetic development of resistance to kinase inhibitors [23]. We have proposed that blocking this adaptive reprogramming is important clinically for making single kinase inhibitors more durable [7]. Pre-and post-drug treatment MIB/MS analysis allows for the quantitative measure of kinome adaptive responses and the rapid screening of combinations of kinase or epigenetic inhibitors that would block the adaptive behavior of the kinome [7,23,24]. This analysis can be done in preclinical models as well as patient trials where biopsy accessible tumor specimens are available. We have been able to capture more than 200 kinases with as little as 300 µg of tumor biopsy protein. Thus, MIBs provide a proteomic approach to characterize the functional state and dynamics of the kinome and thus define therapeutic response and targetable adaptive resistance networks. Importantly, MIBs capture both well-characterized and understudied kinases for a comprehensive measure of the functional kinome.

## Materials and Methods

### MIB affinity chromatography

Broad spectrum Type I kinase inhibitors (CTx-0294885, VI-16832, PP58, Purvalanol B, UNC-2147A, and UNC-8088A) were custom-synthesized with hydrocarbon linkers and terminal amine groups and covalently attached to ECH-activated Sepharose beads as previously described [7]. Cells were rinsed in PBS and processed in lysis buffer (50 mM HEPES, 150 mM NaCl, 0.5% Triton X-100, 1 mM EDTA, 1 mM EGTA, at pH 7.5 containing 10 mM NaF, 2.5 mM NaVO4, cOmplete protease Inhibitor Cocktail (Roche), and 1% Phosphatase Inhibitor Cocktails 2 and 3 (Sigma)). Tumor biopsies obtained from UNC Tissue Procurement were manually homogenized with a chilled mortar and pestle in lysis buffer. For individual bead profiling (Figure 1), 2mg of total protein was gravity-flowed over 100uL of each bead. For Figure 2 (cell lines), 5mg of total protein lysate and for Figure 4 (human tumor biopsies), 1mg of total protein was gravity-flowed over a mixture of the six kinase inhibitor-linked beads (175uL total beads). Beads were washed with at least 30 volumes of high salt (1M NaCl) and low salt (150mM NaCl) lysis buffer, then 500uL of low salt lysis buffer containing 0.1% SDS. Bound proteins were eluted by boiling with 0.5% SDS and 1% β-mercaptoethanol in 100mM Tris-HCl, pH 6.8, 2X 15 minutes, treated with DTT (5mM, 25min at 60°C) and Iodoacetamide (20mM, 30min in the dark at RT), and spin-concentrated to 100µL (Millipore Amicon Ultra-4, 10K cutoff) before Methanol/Chloroform precipitation. Proteins were trypsinized overnight at 37°C and then dried down in a speed-vac. Peptides were cleaned with C-18 spin columns (Pierce).

### Mass Spectrometry and Analysis

Peptides were resuspended in 2% ACN and 0.1% Formic Acid. For Figure 1 (bead profiling) 20% of each sample was injected onto a Thermo Easy-Spray 75µm x 15cm C-18 column using an Easy nLC-1000 in technical triplicate and separated on a 150 min gradient (5-40% ACN). For Figures 2 and 4 (cell lines and tumor biopsies), 40% of the final peptide suspension was injected onto an Easy-Spray 75µm x 25cm C-18 column and separated on a 300min gradient (cell lines) or a 180min gradient (tumor biopsies). For all runs, ESI parameters: 3e6 AGC MS1, 80ms MS1 max inject time, 1e5 AGC MS2, 100ms MS2 max inject time, 20 loop count, 1.8 m/z isolation window, 45s dynamic exclusion. Spectra were searched against the Uniprot/Swiss-Prot database with Sequest HT on Proteome Discoverer software (Figures 1 and 2) or MaxQuant (Figure 4). Only peptides with medium or greater confidence (5% FDR) were considered for quantitation, and only kinases having 3 or more unique peptides were considered for further analysis. Heat maps were generated using GENE-E software (BROAD institute). Kinome trees were generated using Kinome Render (http://bcb.med.usherbrooke.ca/kinomerender.php).

### RNA-seq

Total RNA was spin column purified using RNeasy Plus Mini kit (Qiagen). Library construction was performed at the UNC Lineberger Comprehensive Cancer Center Genomics Core and the sequencing at the UNC High-Throughput Sequencing Facility. mRNA-Seq libraries were TM constructed with 1 µg total RNA using the Illumina TruSeq™ RNA Sample Prep Kit according to the manufacturer’s protocol. 50-cycled single-end sequencing runs with multiplexing were produced using an Illumina HiSeq2000. CASAVA 1.8.2 generated bases and assessed sequence quality. The QC-passed reads were aligned to the hg19 human reference genome using MapSplice and the alignment profile was determined by Picard Tools v1.64 [25]. Aligned reads were sorted and indexed using SAMtools, and then translated to transcriptome coordinates and filtered for indels, large inserts, and zero mapping quality using UBU v1.0. Transcript abundance estimates for each sample were determined using an Expectation-Maximization algorithm [26]. Publicly available data from [27,28] were also processed using this computational method. Data is available in Supplemental Data File S2.

### Data analysis

Hierarchical clustering, Principal Components Analysis (PCA), and feature selection were performed in MATLAB. PCA is a commonly-used data analysis and dimension-reduction technique that transforms variables into a set of linearly uncorrelated principal components [29]. Application of PCA also provides the ability to assign a weight to each feature (kinase) in the data set that can be used as a relative measure of its ability to distinguish subtypes. To identify kinases that dominate individual PCs, kinases having weights in the 90th-percentile (i.e. those weighted in the top 10% of weights) per PC were selected from the first three PCs and used in downstream classification tasks. Feature selection using the Bhattacharyya distance was also used as a secondary mechanism for ranking kinases in terms of their ability to distinguish subtypes [22]. Pairwise classification between subtypes (e.g. basal-like subtype from all others, claudin-low from all others, etc) was iteratively performed to identify the most informative features.

Kinases identified through feature ranking and PCA are combined to create a list of the most distinguishing kinases in MIB-binding across the breast cancer subtypes. Subtype-specific signature kinases are defined as the top 5% of the highest ranking kinases found using the Bhattacharyya feature ranking coefficient for each subtype are compiled for the overall list. Pan-subtype signature kinases are defined as the most heavily weighted kinases (top 10%) from the first three PCs are used. Subtype-specific signature kinases are compiled from each of the breast cancer subtypes then the global signature kinases are added (in order from most heavily weighted to less heavily weighted) starting with PC1 kinases then moving to PC2 then to PC3 until a maximum of 50 kinases is reached to make up the list of distinguishing kinases. A total of 50 kinases was chosen as classification accuracy of the breast cancer subtypes plateaued at this level (Figure S1).

### Comparison of MIB-binding to transcript abundance

The Z-score is calculated by sample based on the average log2 value per kinase and using the standard deviation of all kinases for a given sample for both data types, MIB/MS and RNA-seq (Figure 3A). The Pearson correlation of individual kinases is calculated for each kinase across the 15 cell lines (not distinguished by subtype) between MIB-binding and RNA transcript levels. The distribution of these correlations is shown in Figure 3B. Figure 3C portrays the kinases detected by MIB/MS, RNA-seq, and both data types for the basal-like cell line HCC1806. Example RNA-seq and MIB/MS levels for kinases with good correlation (>0.8) between the data types (ERBB2 and PDPK1) and kinases with poorer correlation (EPHA1 and M3KL4).

### Prediction of subtypes

Classification of subtype based on a previously unobserved kinome signature was performed using a Support Vector Machine (SVM). The SVM is a commonly used machine learning technique used in supervised classification, and thus requires a training set on which to learn parameters that can then be applied towards prediction of previously unobserved data [30]. The SVM used here was trained on the 50 distinguishing kinases previously identified. Performance of the SVM was analyzed using leave-one-out cross validation, where training is performed on all samples except for one and a classification prediction being carried out for the left-out sample. Predictions are made in this way for every sample and final sensitivity, specificity, and precision are calculated on classification performance across all samples. Human tumors are classified into one of the major groups (TNBC or HER2+/Luminal) or as “other” using the SVM trained on the 50 distinguishing kinases identified from breast cancer cell line samples.

### Network Analysis

Protein-protein interaction information was compiled from multiple public data sources for the 254 kinases analyzed in this data set and included, the Human Integrated Protein-Protein Interaction rEference (HIPPIE) (updated 9/1/2015; [31]), Human Protein Reference Database (HPRD Release 9; [32]), Interlogous Interaction Database (I2D version 2.9; [33,34]), PhosphoSitePlus (phosphosite.org - downloaded 10/15/2015; [35]) and Reactome protein-protein interactions (downloaded 12/15/2015; [36]). The union of all interactions between the 254 kinases was used to form a single network that was then clustered into communities/subnetworks with the spectral method in Mathematica (ver 10.3). Subnetworks were assessed for GO-term enrichment via Panther [37]

### Annotation of Tdark Kinases

To identify sets of kinases that were functionally linked through common MIB-binding behavior, we utilized Lasso regression, which has strong utility as a feature selection tool [38]. Lasso regression was performed on each of the 254 kinases which passed initial filtering (from the Supplemental Data File S1). Iterating through all kinases, a single kinase’s MIB/MS data was used as the response vector while all other kinases formed the input matrix. The regression was performed in R utilizing the glmnet package [39]. The resulting features for each Tdark kinase, and the kinases which had it as a resulting feature, are all labeled as regression correlations in Figure 5. Known interactions from earlier described public data sources were identified for each Tdark kinase and each of its regression correlations. The final Tdark kinase networks were created from the combination of both regression correlations and known interactions. All kinases involved in each Tdark kinase network were listed and compared against a list of all human kinases to find overrepresented GO biological processes, Kegg and Reactome pathways via Panther [37] and g:Profiler [40].

## Supplementary Materials

Fig. S1. Classification accuracy across subtypes in LOOCV.

Table S1. Kinases bound to individual beads.

Table S2. Understudied kinases list.

Table S3. Precision, specificity, and sensitivity for all four subtypes in SVM.

Table S4. Kinases bound uniquely within a subtype.

Table S5. Kinases within subnetworks.

Data file S1. Baseline MIB-binding data matrix.

Data file S2. Matrix of RSEM values across cell lines.

Data file S3. MIB-binding response data matrix.

Data file S4. Unique kinases by subtype.

Data file S5. Kinase subnetworks.

## Author contributions

KALC did computational analysis, generated figures, writing, and editing. TS generated experimental (MIB/MS) data, created figures, writing, and editing. JSZ generated experimental (RNA-seq), writing, and editing. MPE did writing and editing. TP generated experimental (MIB/MS) data. CRH generated growth curves, functional annotation, writing, and editing. DRG generated experimental (MIB/MS) data, writing, and editing. SMB generated experimental (MIB/MS) data, writing, and editing. SPA generated experimental (MIB/MS) data, writing, and editing. SHV generated experimental (MIB/MS) data, writing, and editing. NS analyzed RNA-seq data in pipeline, writing, and editing. TIO contributed to functional annotation. LMG was involved in experimental design. GLJ managed and designed experimental work, writing, and editing. SMG managed and designed computational analyses, writing, and editing.

## Data and materials availability

All processed data is provided in Supplementary Data Files. Data from the Sequence Read Archive is used for some analyses here: SRX317702, SRX317711, SRX317712, SRX317715, SRX317717, SRX317723, SRX317730, SRX317733, SRX317736, SRX317738PT1, SRX317738PT2, SRX317741, SRX317743, SRX317747.

